# Integrative proteomic and phosphoproteomic analysis of granulosa cells during follicular atresia in porcine

**DOI:** 10.1101/2020.08.31.276436

**Authors:** Feng Yang, Qiang Liu, Yanhong Chen, Huizhen Ye, Han Wang, Shenming Zeng

## Abstract

Ovarian follicular atresia is a natural physiological process; however, the mechanism is not fully understood. In this study, quantitative proteomic and phosphoproteomic analyses of granulosa cells (GC) in healthy (H), slightly atretic (SA), and atretic follicles (A) of porcine were performed by TMT labeling, enrichment of phosphopeptides and LC-MS/MS analysis. In total, 6,201 proteins were quantified and 4,723 phosphorylation sites of 1,760 proteins were quantified. In total, 24 (11 up, 13 down) and 50 (29 up, 21 down) proteins with a fold change (FC) > 5 were identified in H/SA and H/A, respectively. In addition, there were 20 (H/SA, up) and 39 (H/A, up) phosphosites with an FC > 7, that could serve as potential biomarkers for distinguishing different quality categories of follicles. Western blotting and immunofluorescence confirmed the reliability of the proteomic analysis. Some key proteins (e.g., MIF, beta catenin, integrin β2), phosphosites (e.g., S76 of caspase6, S22 and S636 of lamin A/C), pathways (e.g., apoptosis, regulation of actin cytoskeleton pathway), transcription factors (e.g., STAT5A, FOXO1, and BCLAF1), and kinases (e.g., PBK, CDK5, CDK12, AKT3) involved in atresia process were revealed via further analysis of the differentially expressed proteins (DEPs) and phosphorylated proteins (DEPPs). Collectively, the proteomic and phosphoproteomic profiling and functional research in the current study comprehensively analyzed the dynamic changes in protein expression and phosphorylation during follicular atresia and provided some new explanations regarding the regulation of this process.

## 1. Introduction

In female mammals, less than 1% of follicles are selected and the remainder are eliminated (1, 2). Most follicles undergo a degenerative process known as atresia (3). Follicular atresia was described as being common among all vertebrate groups in early 1947 (4, 5). In humans, the total germ cell number reaches a peak of 6.8 million at 5 months of gestational age. By the time of birth, this number has declined to 2 million and only 300–400 thousand follicles are present at the onset of puberty (6, 7). A woman usually ovulates only approximately 400 follicles during her reproductive life (7); therefore, more than 99.9% of human follicles undergo atresia. In porcine, the reserve of primordial follicles is estimated to be 10 million in the ovaries 10 days after birth and the recruitment of resting primordial follicles into the growing pool begins during the fetal life (3). However, at most, 1600 oocytes will ovulate during the fertile life in porcine, and the others will disappear (3). Atresia occurs continually throughout a female's life and follicular atresia is similar in pigs and humans. Pigs are economically important and have also been increasingly used as an alternative model for studying human health and disease and for testing new surgical and pharmacological treatments (8). In addition, pigs are closely related to humans in terms of anatomy, genetics and physiology (9). Researchers often use porcine follicles to study follicular development and atresia, because there are various grades of follicles on their ovaries, which are easy to observe and manipulate.

It is widely accepted that ovarian follicular atresia is mediated primarily by granulosa cell (GC) apoptosis (10–13), which is regulated by internal and external factors (14). The GCs first undergo apoptosis, then the GC layer detaches from the follicular basement membrane (BM), after which, fragmentation of the BM begins and finally, the follicle disappears (3). Apoptosis of GCs is similar to other cell types, which exhibit cell membrane blebbing, DNA degradation, and protease activation; however, there are some specific characteristics of GC apoptosis. For example, during the initial steps of apoptosis, steroidogenesis is increased due to aggregation of the steroidogenic organelles in the perinuclear region and their exclusion from the apoptotic blebs. Actin cytoskeleton reorganization plays an important role in this compartmentalization as well as in transmitting survival factors exerted by the BM (15). Atresia is triggered when some essential factors supporting follicular development are lacking. In particular, the gonadotropin (FSH and LH), growth factors, cytokines, steroids, and constituents of extracellular matrix (ECM) (13). The Bcl-2 protein family plays an irreplaceable role during apoptosis. Recent results from experiments done by our group demonstrated that heat stress promotes BimEL phosphorylation at Thr112 through the JNK pathway and decreases the level of aromatase in porcine GC, thus damaging follicular development (16, 17).

To study the dynamic changes in gene expression during the process of follicular atresia, transcriptome analysis was performed in porcine GCs from healthy and atresia follicles by two independent laboratories (18, 19). Some markers of follicular atresia, key genes, and pathways were found; however, the ultimate function executor in biological processes is proteins. The continuous advancement of proteomics technology provides the opportunity to gain a deeper understanding of the dynamic expression profiles of proteins during atresia. Protein phosphorylation is one of the most important posttranslational modifications (PTM) in living cells and is involved in various biological processes, including follicular atresia (16, 17). However, the dynamic spectrum of proteins and protein phosphorylation during the process of follicular atresia has not yet been elucidated. In addition, dynamics of transcription factors (TFs) and kinases during this process are not yet known. In this study, quantitative proteomic and phosphoproteomic analyses of the GCs of healthy (H), slightly atretic (SA), and atretic (A) porcine follicles were performed to further investigate the mechanisms of follicular atresia.

## 2. Materials and Methods

### 2.1 Classification of Healthy, Slightly Atretic, and Atretic Follicles and Recovery of GCs

Ovaries from gilts aged about 5 months were collected at a local abattoir and transported to the laboratory in a vacuum flask containing sterile physiological saline (30–35 □) within 2 h of collection. Ovaries were washed twice with sterile physiological saline (37 □) containing 100 IU/L penicillin and 50 mg/L streptomycin. H, SA, and A follicles in porcine were classified according to previously established morphological criteria (18, 20, 21). Follicular (3–5 mm in diameter) contents from 10 ovaries were punctured by hypodermic needle, and cumulus-oocyte complex and ovarian tissue were discarded under a stereo microscope. The GCs were subsequently harvested by centrifuging.

### 2.2. Protein Extraction, Concentration Measurement, and Trypsin Digestion

Protein extraction, concentration measurement, and trypsin digestion were done as previously described (22). Cells were lysed in lysis buffer containing 8 M urea, 10 mM DTT, 1% Protease Inhibitor Cocktail and phosphatase inhibitors. Samples were centrifuged to remove debris, and the supernatant was collected. Finally, the protein was precipitated with cold 15% TCA for 2 h at 4 °C. After centrifugation at 4 °C at 5000 g for 10 min, the supernatant was discarded. The remaining precipitate was washed three times with cold acetone. The protein was then dissolved in buffer (8 M urea, 100 mM TEAB, pH 8.0) and the protein concentration was determined using a 2-D Quant kit (GE Healthcare, Piscataway NJ, USA) according to the manufacturer’s instructions. For digestion, the protein solution was reduced with 10 mM DTT for 1 h at 37 °C and alkylated with 20 mM IAA for 45 min in the dark at room temperature. For trypsin digestion, the protein sample was diluted by adding 100 mM TEAB to urea concentration less than 2M. Finally, trypsin was added at a 1:50 trypsin-to-protein mass ratio for the first overnight digestion and at a 1:100 trypsin-to-protein mass ratio for the second 4 h-digestion.

### 2.3. TMT Labeling, HPLC Fractionation and Affinity Enrichment of Phosphopeptides

After trypsin digestion, peptide was desalted by Strata X C18 SPE column (Phenomenex, California, USA) and vacuum-dried. Peptide was reconstituted in 1 M TEAB and processed using a 6-plex TMT kit according to the manufacturer’s protocol. Briefly, one unit of TMT reagent (defined as the amount of reagent required to label 100 μg of protein) was thawed and reconstituted in 24 μl ACN. The peptide mixtures were then incubated for 2 h at room temperature, pooled, desalted, and dried by vacuum centrifugation.

HPLC fractionation and affinity enrichment of phosphopeptides were done as previously described (22). Briefly, the peptides were combined into 8 fractions and dried by vacuum centrifuging. The phosphopeptides were enriched using IMAC (immobilized metal affinity chromatography). The supernatant containing phosphopeptides was collected and lyophilized for LC-MS/MS analysis.

### 2.4. LC-MS/MS Analysis

LC-MS/MS analysis was performed as previously described (22, 23). Briefly, after the peptides were dissolved using 0.1% formic acid, samples were separated using an EASY-nLC 1000 ultrahigh performance liquid phase system, and then subjected to NSI source followed by Q-Exactive Plus system analysis. Peptides were selected and fragmented with a normalized collision energy of 30 eV fragment ions were detected in the Orbitrap at resolution of 17,500. For the proteomic. The gradient was comprised of an increase from 7% to 25% solvent B (0.1% FA in 98% ACN) for 24 min, 25% to 40% for 8 min and climbing to 80% in 4 min then holding at 80% for the last 4 min, all at a constant flow rate of 350 nl/min on an EASY-nLC 1000 UPLC system. For the phosphorproteomic. The gradient was comprised of an increase from 4% to 22% solvent B (0.1% FA in 98% ACN) for 40 min, 22% to 35% for 12 min and climbing to 80% in 4 min then holding at 80% for the last 4 min, all at a constant flow rate of 400 nl/min on an EASY-nLC 1000 UPLC system.

### 2.5. Proteomic and phosphorproteomic database search

Raw data were processed with Max MaxQuant search engine (v.1.5.2.8). Tandem mass spectra were searched against uniprot *sus scrofa* (26201 sequences) database. For the proteomics, Trypsin/P was specified as the cleavage enzyme allowing up to 2 missing cleavages. Mass error was set to 20 ppm for precursor ions and 0.02 Da for fragment ions. Carbamidomethyl on Cys was specified as fixed modification, oxidation on Met and Acetylation on protein N-term were specified as variable modifications. For the protein quantification method, TMT 6-plex was selected in Maxquant. The FDR was adjusted to < 1% at the protein and PSM level. For the phosphoproteomics, Trypsin/P was specified as the cleavage enzyme allowing up to 2 missing cleavages. Maximun number of modifications per peptide was setting 5. Mass error was set to 20 ppm for precursor ions and 0.02 Da for fragment ions. Carbamidomethylation on Cys was specified as fixed modification and oxidation on Met, phosphorylation on Ser, Thr, Tyr, and acetylation on protein N-terminal were specified as variable modifications. False discovery rate (FDR) thresholds for protein, peptide, and modification site were specified at 1%. Minimum peptide length was set at 7. For the quantification method, TMT-6-plex was selected. The site localization probability was set as > 0.75. The mass spectrometry proteomics data have been deposited to the ProteomeXchange Consortium via the PRIDE partner repository with the dataset identifier PXD020899.

### 2.6. GO, domain, KEGG pathway annotation and subcellular localization

UniProt-GOA database was used to annotate protein GO terms. Proteins domain were annotated by InterProScan based on protein sequence alignment method, and the InterPro domain database was used. Kyoto Encyclopedia of Genes and Genomes (KEGG) database was used to annotate protein pathway. We used wolfpsort, a subcellular localization predication soft to predict subcellular localization.

### 2.7. Motif analysis

Soft motif-x was used to analysis the model of sequences constituted with amino acids in specific positions of modify-13-mers (6 amino acids upstream and downstream of the site) in all protein sequences. And all the database protein sequences were used as background database parameter, other parameters with default.

### 2.8. Protein functional enrichment

For each protein functional term, a two-tailed Fisher’s exact test was employed to test the enrichment of the differentially expressed protein against all identified proteins. Term with a corrected *P* < 0.05 were considered significant.

### 2.9. Functional enrichment-based clustering

*P* value of functional enrichment was collated from compare groups. retaining functional term which was at least enriched in one of the compare group with *P* <0.05. Then P value matrix was transformed by −log10 and z-score for each functional category. These z scores were clustered by one-way hierarchical clustering (Euclidean distance, average linkage clustering). Cluster membership were visualized by a heat map using the “heatmap.2” function from the “gplots” R-package.

### 2.10. Transcription factor searching

The 1,490 known porcine TFs from the Animal Transcription Factor Database (AnimalTFDB3, http://bioinfo.life.hust.edu.cn/AnimalTFDB/) were used to search and analyze the expression patterns of the TFs in the proteome and phosphoproteome (24).

### 2.11. Kinase analysis

GPS 5.0 software was used for predicting kinase-substrate regulations. The corresponding kinase proteins in the kinase family were obtained by comparison with the kinase sequence in the IEKPD2.0 database. Protein-protein interaction (PPI) information was used to filtrate potentially false-positive hits. A “medium” threshold was chosen in GPS 5.0. The GSEA method was used to predict kinase activities, in which log-transformed phosphorylation levels (or ratio values) as a rank file and kinase-phosphorylation site regulations were formatted into a GMT format file in a sample (or a comparable group). Normalized enrichment scores (NES) were obtained from enrichment result and were regarded as kinase activity scores. Kinase were predicted as positive if the predominant change in substrates was an increase in phosphorylation and visa-versa. For each comparable group, kinases predicted as having positive or negative activity and as having significantly differentially expressed phosphorylation sites were used to construct kinase-substrate regulatory network, according to the complicated regulatory relationships.

### 2.12. Protein Extraction and Immunoblotting

The GCs were harvested and washed once in PBS, then lysed on ice for 30 min with RIPA buffer (CST, 9806), and supplemented with 1% (v/v) protease inhibitor Cocktail (HY-K0010) and 1% (v/v) phosphatase inhibitors (Cocktail I, HY-K0021; Cocktail II, HY-K0022; and Cocktail III, HY-K0023), which were purchased from MCE (Shanghai, China). Western blotting was performed as described previously (16, 17). Protein concentrations were determined using a BCA protein assay kit (Transgen Biotech, Beijing, China). Equal amounts of proteins (15–50 μg/lane) were separated by SDS-PAGE (12% acrylamide running gel) and transferred to a nitrocellulose membrane (BioTrace™ NT, Pall Corp, FL, USA). The following antibodies were used in this experiment: beta catenin (ab32572, Abcam), inhibin alpha (ab81234, Abcam), HSD17B1 (ab134193, Abcam), MIF (ab227073, Abcam), caspase6 (ab185645, Abcam), laminA/C (MA3-1000, Thermo), p-laminA/C-S22 (13448, CST, Shanghai, China). The antibodies were diluted to the recommended ratio with Beyotime (P0256, Shanghai, China) diluent. The Western blotting images were analyzed using Image J software (National Institutes of Health, Bethesda, MD, USA).

### 2.13. Immunohistochemical and immunofluorescence

Porcine follicles (3–5 mm in diameter) of different health statuses were fixed overnight in 4% phosphate-buffered formaldehyde at 4 °C for 7 days and then embedded in paraffin. Randomly selected sections (5 μm each) were used for subsequent staining. Immunohistochemical and immunofluorescence were performed as previously described (16, 17). MIF (1:500), laminA/C (1:100), cacpase6 (1:1000) antibodies were used for staining. Immunohistochemical slides were examined under light microscopy (Leica DC200 digital camera; Leica, Germany). Immunofluorescence samples were examined using a confocal microscope (A1HD25, Nikon, Japan), and images were recorded.

### 2.14. Detection of the MIF concentration in follicular fluid and MIF mRNA relative expression in GCs

Follicular (3–5 mm in diameter) contents from H, SA, and A follicles were punctured by hypodermic needle. Follicle fluid was obtained via centrifuging at 600 g for 10 min. MIF concentration in the follicular fluid was detected using a MIF ELISA kit (JLC30201, Shanghai Jichun Industrial Co., Ltd. China). MIF mRNA relative expression in GCs was detected via Q-PCR. MIF primer: forward: ATCAGCCCGGACAGGATCTA, reverse: GCCGAGAGCAAAGGAGTCTT.

### 2.15. Statistical Analyses

Proteins, MIF concentration, and MIF mRNA relative expression levels (gray values) are presented as the means ± SDs and were analyzed using a one-way ANOVA with SPSS 22 (IBM, SPSS, Chicago, IL, USA) for Windows. ANOVA followed by a post hoc Dunnett’s test was used to determine differences between different groups. Differences were considered statistically significant at *P* < 0.05.

## 3. Results

### 3.1. Overview of proteomic and phosphoproteomic profiling of porcine GCs during atresia

The proteomic and phosphoproteomic analysis workflow is illustrated in Fig. 1A. The GC samples from H, SA, and A were lysed, digested, and labeled with different TMT tags, then pooled and analyzed by LC/LC-MS/MS. Of the pool, 5% of the pool was used for proteome analysis, and the remaining 95% was subjected to phosphoproteome profiling. Altogether, 6,201 of the 7,104 identified proteins were quantified. In addition, 6,839 phosphorylation sites in 2,830 proteins were identified, among which 4,723 phosphorylation sites in 1,760 proteins were quantified. The length of most peptides varied between 8 and 20 in the proteomic and phosphoproteomic data, indicating that the sample preparation reached the standard (Fig. 1B). Reproducibility analysis indicated high reproducibility between biological duplicate samples (Fig. 1C). Relative quantitation of proteins was divided into two categories: a quantitative ratio over 1.5 was considered upregulation while a quantitative ratio less than 0.67 was considered downregulation. The amount of differentially expressed proteins (DEPs) in GC from different comparable groups are summarized in Fig. 1D. The amount of the differentially quantified phosphorylation sites and differentially expressed phosphorylated proteins (DEPPs) from different comparable groups are summarized in Fig. 1E.

**Fig. 1.**
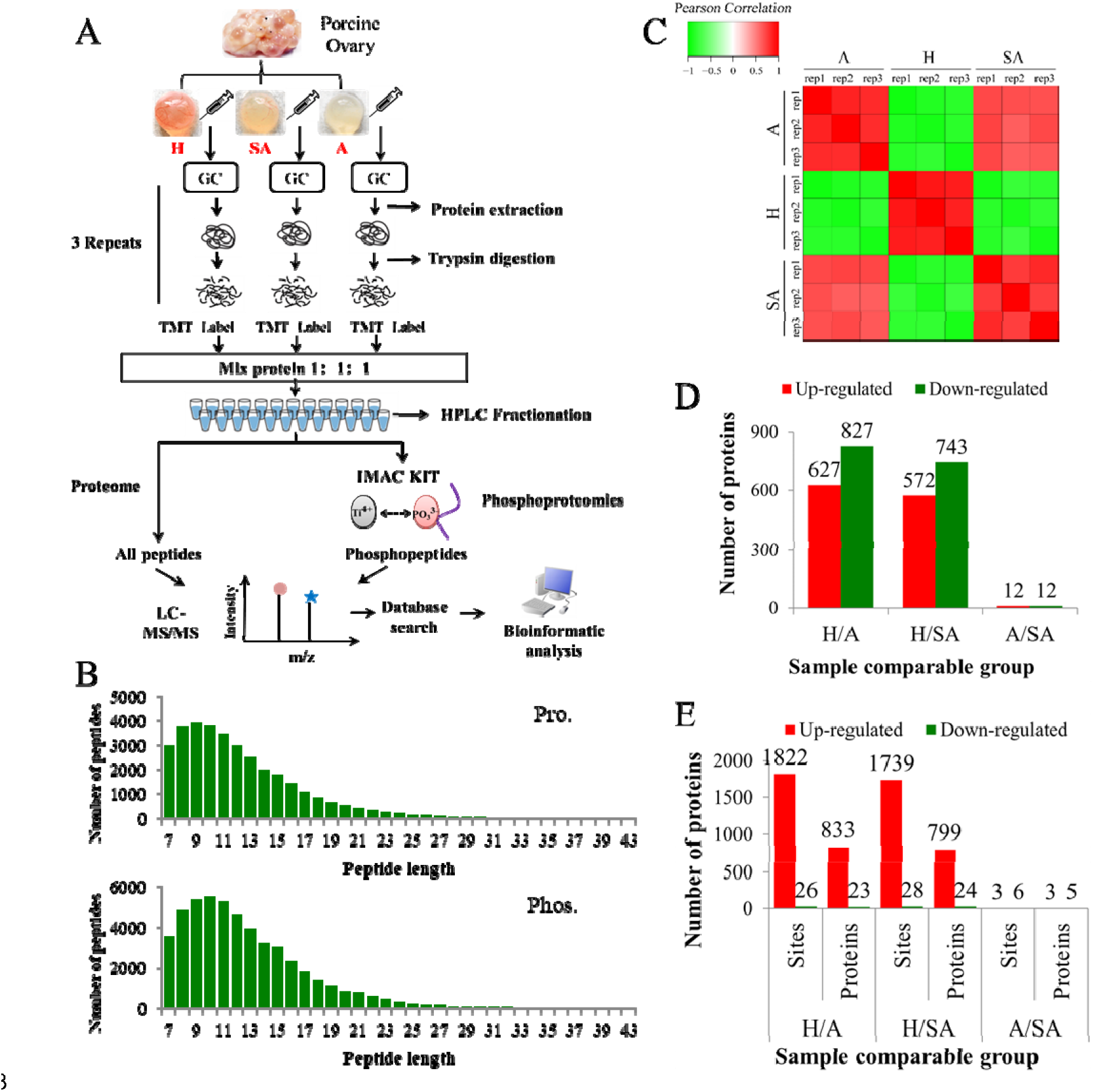
Overview of proteomic and phosphoproteomic analysis in granulosa cells obtained from ovaries with different follicular statuses. A. Proteomic and phosphoproteomic analysis workflow in granulosa cells from healthy (H), slightly atretic (SA) and atretic (A) porcine follicle samples. B. The distributions of raw MS/MS length of peptides and phosphopeptides quantified from proteomic and phosphoproteomic data, respectively. C. Reproducibility analysis of the samples. D. The amounts of DEPs in different comparable groups. E. The amounts of the differentially quantified phosphosites and phosphoproteins in different comparable groups.

### 3.2. The global proteome changs in GCs during porcine follicular atresia

In this study, 6,201 of 7,104 identified proteins were quantified. These data are presented in Table S1 and the DEPs are presented in Table S2. Among these, 24 (11 up, 13 down) proteins in H/SA, and 50 (29 up, 21 down) proteins in H/A had a fold change (FC) higher than five, which could be used as potential novel biomarkers to distinguish different health statuses of follicles, especially between H and SA. DEPs in H/SA follicles with a FC ≥ 5 are shown in Table 1.

**Table 1.**
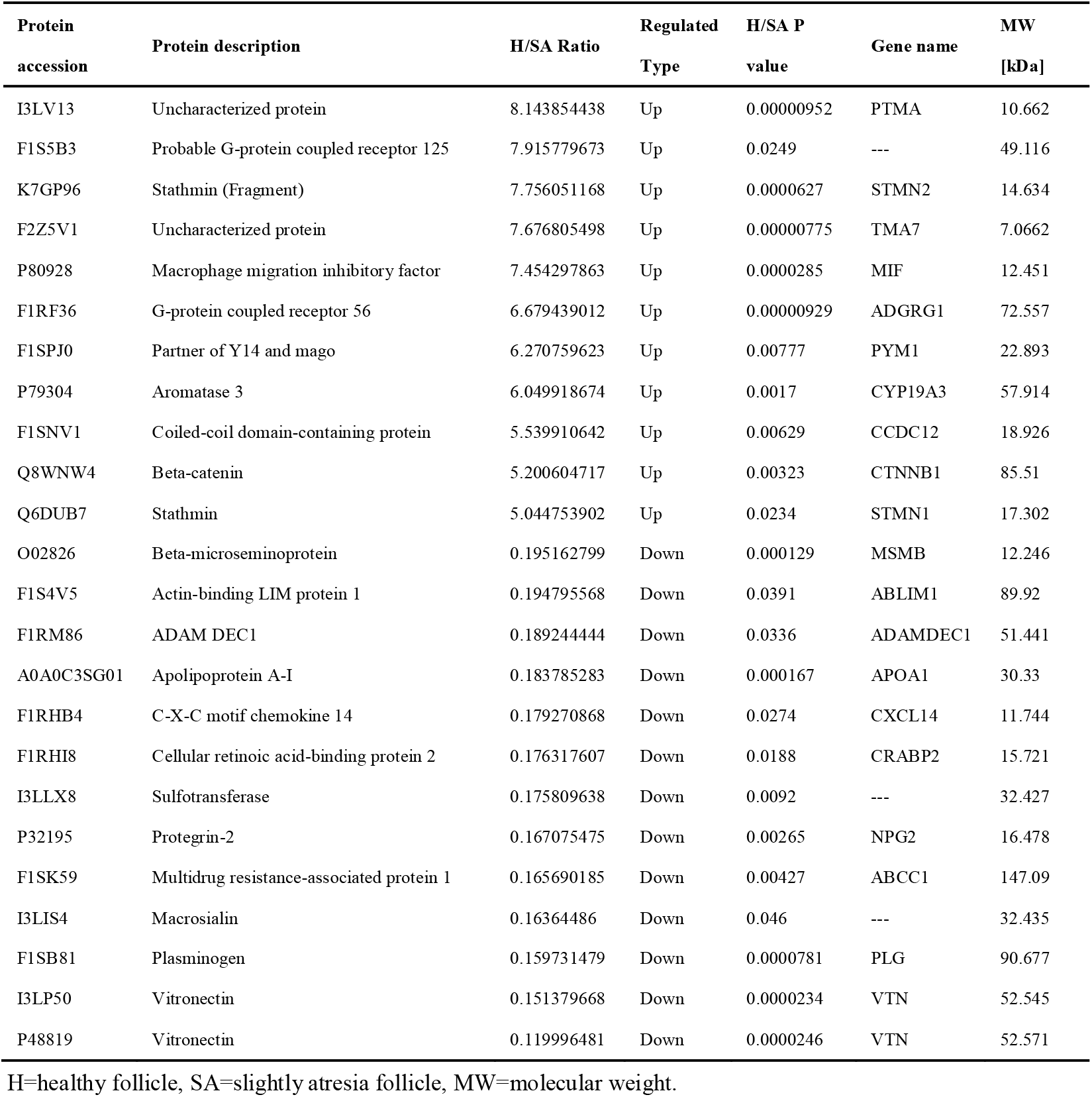
DEPs in H/SA follicles with a FC higher than 5.

Five randomly selected DEPs (β-catenin, laminA/C, inhibin alpha, HSD17B1, and MIF) were validated via Western blotting, thus proving the reliability of the proteomic data (Fig. 2A). MIF was 7.45 times higher in GCs from H compared to SA follicles, and 13.57 times higher in GCs from H compared to A follicles and was confirmed by Western blotting and immunohistochemistry (Fig. 2A and 2B). Immunofluorescence results showed that MIF was primarily expressed in GCs of primordial, primary, secondary, and small antrum follicles, and was also expressed in cumulus cells (Fig. 2C and 2D). MIF concentration in follicular fluid was highest in H compared with SA and A follicles (Fig. 2E). In addition, the MIF mRNA expression level in GCs was the same as the protein profile in all three follicular categories (Fig. 2F), which is consistent with a previous transcriptome study of porcine follicular atresia (18). MIF can function as cytokine, hormone, and enzyme; however, its physiologic significance has yet to be elucidated (25, 26). In addition to inflammatory and immunologic functions, MIF plays a role in tumor cell growth, cell proliferation, would repair, regulation of cytochrome c release, and inhibition of Bim-induced apoptosis (27–30). Earlier studies in humans and mice revealed that MIF affects GC and theca cell proliferation, follicular growth, and ovulation (31, 32). MIF was discovered in 1966 and is a molecule capable of inhibiting the random migration of macrophages (33). Ovarian macrophages are important regulators of the complex communication between the immune and reproductive systems (34). Macrophages enter the ovarian follicle at the time of initiation of GC apoptosis, and migrate as apoptosis processes (35). Macrophages can modulate follicle development by secreting EGF and other cytokines and suppress follicular cell apoptosis (36). Macrophages may facilitate angiogenesis, express growth factors to enhance follicle growth, and secrete proteases to remodel the ECM (37). The IL-33-regulated autophagy-macrophage mechanism is required for disposal of degenerative tissues in ovaries in order to maintain ovarian tissue integrity and to retard the aging process (38). Taken together, this implies that MIF may play important role in follicular development and could be used as a novel biomarker to distinguish different categories of follicle quality.

**Fig 2.**
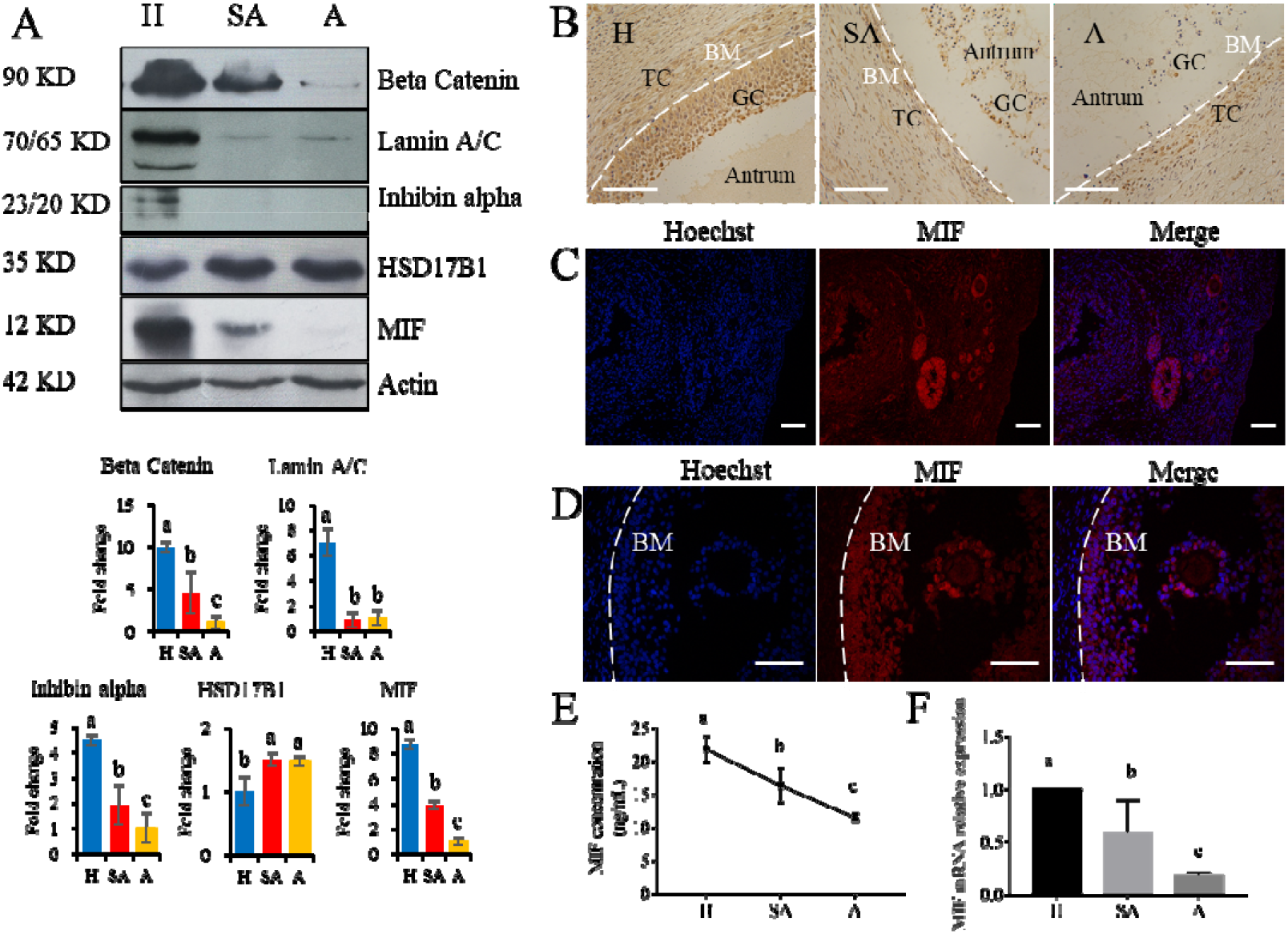
Western blotting validation of expression levels of target proteins found differentially expressed among H, SA, and A follicles by proteomics. A. Protein levels of beta catenin, laminA/C, inhibin alpha, HSD17B1, and MIF were detected by Western blotting in granulosa cells from healthy, slightly atretic and atretic porcine follicles. The histogram shows the quantitative analysis (mean ± SD) of the results of three independent Western blots. The bars are labeled with completely different letters (a, b, c) indicating significant difference, *P* < 0.05. B. Immunohistochemical detection of MIF expression in pig healthy, slightly atretic, and atretic porcine follicles. C. MIF is primarily expressed in granulosa cells of primordial follicles, primary follicles, secondary follicles and small antrum follicles. D. MIF is also expressed in cumulus cells. E. MIF concentrations in follicular fluid from H, SA, and A follicles. F. MIF mRNA levels in GCs from H, SA, and A follicles. H: healthy follicle; SA: slightly atretic follicle; A: atretic follicle; BM: basement membrane; GC: granulosa cell; TC: theca cell. Scale bar is 100 μm. The data are representative of at least three independent experiments.

#### Functional classification of DEPs

The amount of the DEPs in each GO term of level 2 were summed up according to Gene Ontology (GO) annotation information of identified proteins; these proteins can be found in S1 File. The results revealed that the DEPs in cellular component were mainly concentrated in cell, organelle, membrane, and macromolecular complex; the DEPs in molecular function were mainly concentrated in binding, catalytic activity, transporter activity, structural molecule activity, molecular function regulator, and nucleic acid binding transcription factor activity; the DEPs in biological process were mainly concentrated in cellular process, metabolic process, single-organism process, biological regulation, response to stimulus, localization, and signaling; and the DEPs in subcellular location were mainly mapped in nucleus, cytoplasm, mitochondria, extracellular, and plasma membrane.

#### Functional enrichment-based cluster analysis

A GO enrichment-based clustering analysis was performed to characterize the functions of the DEPs in GCs of different follicular statuses during atresia, and included biological process, cellular component, KEGG pathway, molecular function, and protein domain (Fig. 3A-3C). In the biological process category (Fig. 3A), amide biosynthetic process and peptide metabolic process were enriched in proteins that are higher in H than in SA; and cell redox homeostasis was enriched among proteins that are higher in H than in A. In contrast, proteolysis and blood coagulation were enriched among proteins that are higher in SA than in H; nucleobase-containing small molecule metabolic process and actin cytoskeleton organization were enriched among proteins that are higher in A than in H. In the molecular function category (Fig. 3B), protein homodimerization activity, identical protein binding, structural molecule activity, and structural constituent of ribosome were upregulated when follicles changed from H to SA. In addition, oxidoreductase activity, isomerase activity, threonine-type peptidase activity, threonine-type endopeptidase activity, and cofactor binding were upregulated when follicle turn from H to A. In contrast, phospholipid binding, phosphatidylinositol binding, GTPase activity, endopeptidase activity, lipid binding, serine-type endopeptidase activity, peptidase activity-acting on L-amino acid peptidase, peptidase activity, and ligase activity were downregulated when follicle turn from H to SA; whereas, enzyme regulator activity, Ran GTPase binding, GTPase binding, Ras GTPase binding, and small GTPase binding were downregulated when follicles changed from H to A. In the cellular component-based clustering analysis (Fig. 3C), non-membrane-bounded organelle, intracellular non-membrane-bounded organelle, intracellular ribonucleoprotein complex, ribonucleoprotein complex, ribosome, peptidase complex, endopeptidase complex, and proteasome complex were up-regulated when follicle turn from H to SA. Proteasome core complex, prefoldin complex, spliceosomal snRNP complex, spliceosomal snRNP complex, proteasome core complex, alpha-subunit complex, and U1 snRNP were upregulated when follicle turn from H to SA and A. In contrast, extracellular region and extracellular region part were downregulated when follicle turn from H to SA and A. Microtubule and extracellular space were downregulated when follicle turn from H to SA. Enrichment-based clustering analyses were performed using the KEGG database to profile the cellular pathways (Fig. 3D). The ribosome, peroxisome and glycolysis/gluconeogenesis pathways were upregulated in H compared with SA. The ECM-receptor interaction as well as cysteine and methionine metabolism pathways were up-regulated when follicles changed from H to A. In contrast, complement and coagulation cascades, AGE-RAGE signaling pathway in diabetic complications, influenza A, fluid shear stress and atherosclerosis, staphylococcus aureus infection, and proteasome were downregulated in H compared with SA. Regulation of actin cytoskeleton, salmonella infection, NF-kappa B signaling pathway, aminoacyl-tRNA biosynthesis, and Epstein-Barr virus infection were down-regulated when follicles changed from H to A. According to the analysis of the protein domain, high mobility group box domain and zona pellucida domain were upregulated when follicle turn from H to SA (Fig. 3E). Thioredoxin-like fold was upregulated when follicles changed from H to A. In contrast, glutathione S-transferase-C-terminal, AGC-kinase-C-terminal, phox homologous domain, tubulin/FtsZ-GTPase domain, tubulin-C-terminal, tubulin/FtsZ-2-layer sandwich domain, and tubulin/FtsZ-C-terminal were downregulated when follicle turn from H to SA. Whereas, rossmann-like alpha/beta/alpha sandwich fold, kringle-like fold, NAD(P)-binding domain, importin-beta-N-terminal domain, p53-like transcription factor-DNA-binding, pleckstrin homology domain, phosphoribosyl transferase-like, zinc finger-LIM-type, armadillo-like helical, and armadillo−type fold were downregulated when follicles changed from H to A.

Enrichment pathway images of DEPs in GCs of different follicular statuses can be found in S2 File.

**Fig 3.**
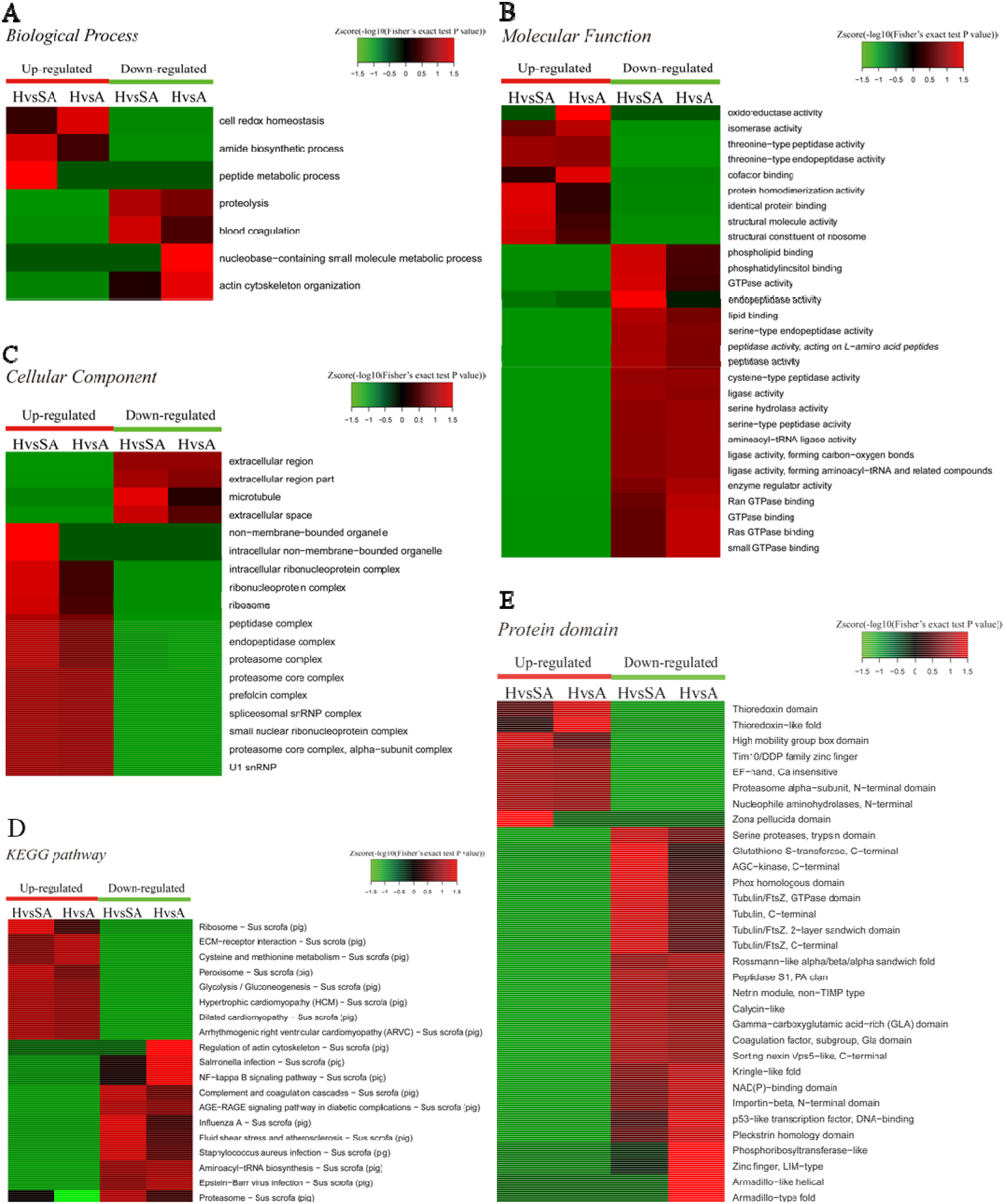
Heat map obtained from functional enrichment-based cluster analysis of the quantified proteomics datasets of different follicular statuses. A, B, and C. Heat maps obtained from GO enrichment-based cluster analysis. D and E. Heat map obtained from enrichment-based cluster analysis of protein domains and KEGG pathways. H=healthy follicle; SA=slightly atretic follicle; A=atretic follicle.

### 3.3. The phosphoproteome changs in GCs during porcine follicular atresia

After IMAC enrichment and the LC-MS/MS analysis, 6,839 phosphorylation sites in 2,830 proteins were identified, and 4,723 phosphorylation sites in 1,760 proteins were quantified. These data are presented in Table S3. The phosphoproteomic data were then normalized to the results of the global proteome analysis. The DEPPs can be found in Table S4. There were 20 (up) and 39 (up) phosphorylation sites that had a FC ≥ 7 in the H/SA and H/A groups, respectively. In addition, 28 phosphorylation sites were downregulated in H/SA, and 26 phosphorylation sites were downregulated in H/A. The DEPPs in H/SA with a FC ≥ 7 are shown in Table 2.

**Table 2.**
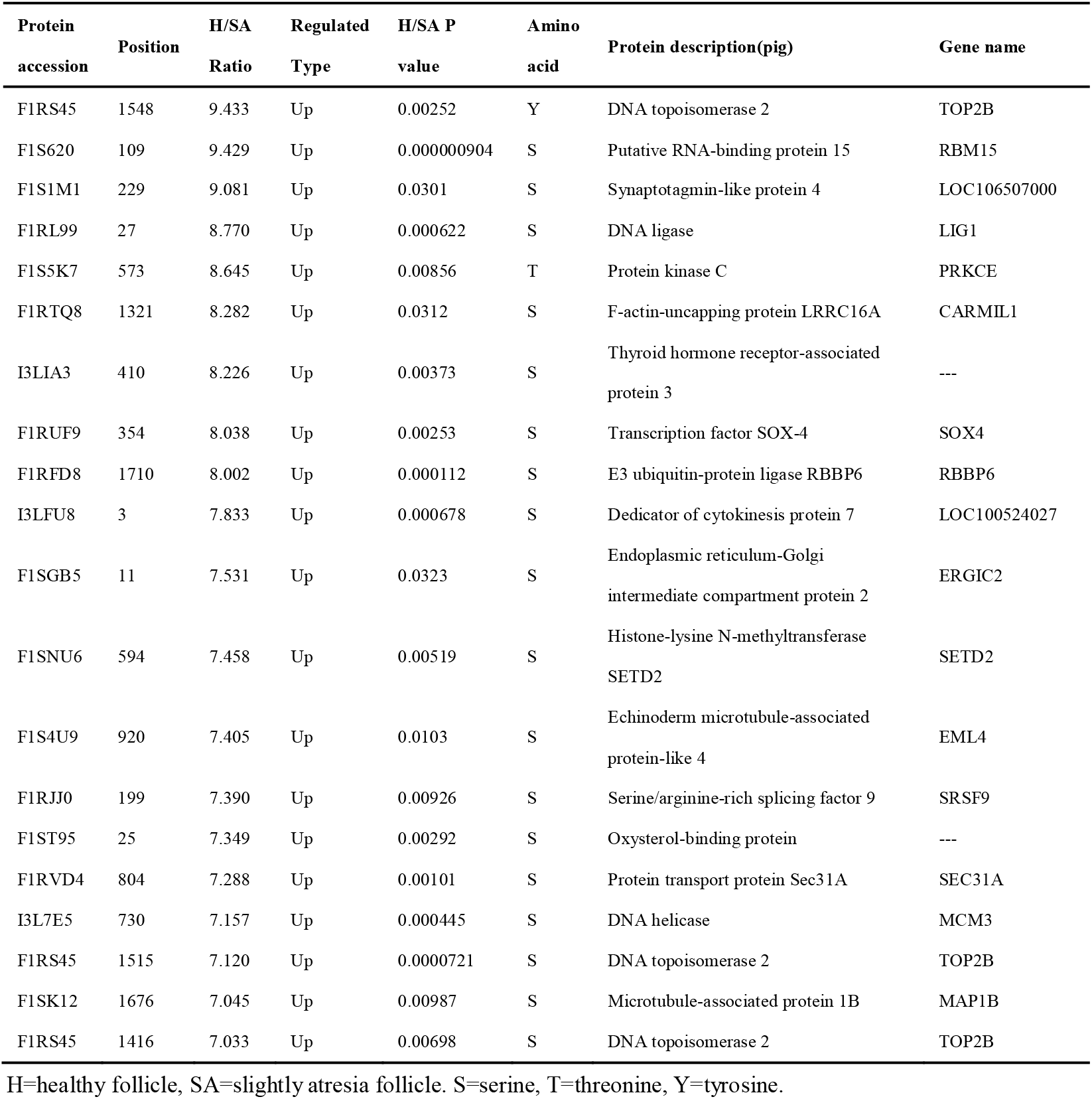
DEPPs in H/SA with a FC greater than 7.

#### Functional classification of DEPPs

The amount of the DEPPs in each GO term of level 2 were summed up according to the GO annotation information of identified proteins; these proteins can be found in S3 File. Similar to the results in the proteome, the DEPPs in cellular component were mainly concentrated in cell, organelle, macromolecular complex, and membrane; the DEPPs in molecular function were mainly concentrated in binding and catalytic activity; and the DEPPs in biological process were mainly concentrated in cellular process, metabolic process, single-organism process, biological regulation, response to stimulus, signaling, and localization.

A functional enrichment-based clustering analysis was performed to characterize the functions of DEPPs in different comparable groups. In the molecular function category (Fig. 4A), DNA topoisomerase activity, transferase activity-transferring one-carbon groups, methyltransferase activity, and RNA binding process were upregulated when follicles changed from H to A. In contrast, structural molecule activity, nucleic acid binding, small molecule binding, nucleotide binding, and nucleoside phosphate binding were downregulated when follicles changed from H to A. In the cellular component category (Fig. 4B), DEPPs were mainly located in the mitochondrial membrane part, mitochondrial protein complex, mitochondrial membrane, mitochondrial outer membrane, organelle outer membrane, cell-cell junction, cell junction, non-membrane-bounded organelle, and intracellular non-membrane-bounded organelle. As shown in Fig. 4C, the biological process analysis revealed that the chromatin organization process was upregulated while the oxoacid metabolic process was downregulated when follicles changed from H to SA. Then, KEGG clustering was performed to characterize the alterations in signaling pathways during follicular atresia (Fig. 4D). Results revealed that mismatch repair, galactose metabolism, spliceosome, vasopressin-regulated water reabsorption, and ubiquitin mediated proteolysis pathways were upregulated when follicles changed from H to SA or A. Importantly, apoptosis pathway was downregulated when follicle turns from H to SA. According to the analysis of the protein domain (Fig. 4E), the PWWP domain and MIF4G-like domain were upregulated when follicles changed from H to SA or A. However, the Lamin Tail domain was downregulated when follicles changed from H to SA, which is involved in the apoptosis pathway.

**Fig 4.**
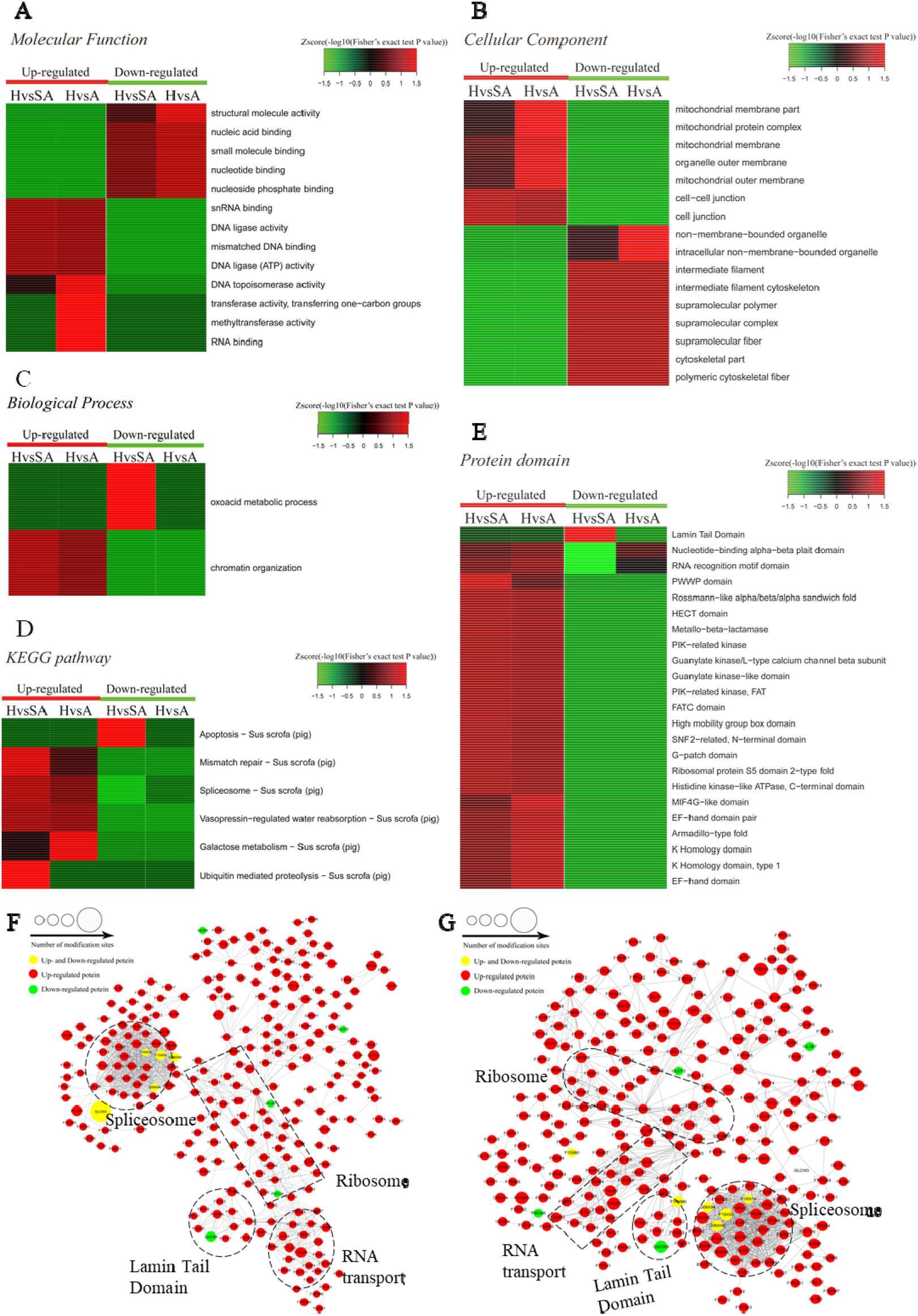
Heat map obtained from functional enrichment-based cluster analysis of quantified phosphoproteome during follicular atresia, and protein-protein interaction networks of DEPs. A, B, and C. Heat maps obtained from GO enrichment-based cluster analysis. D and E. Heat map obtained from enrichment-based cluster analysis of protein domains and KEGG pathways. Analyses of the protein-protein interaction networks for di□erentially expressed phosphorylated proteins between H and SA (F), and between H and A (G). H=healthy follicle; SA= slightly atretic follicle; A=atretic follicle.

Protein-protein interaction networks were established based on the proteins whose phosphosites were regulated during follicular atresia. Molecular complex detection showed that these proteins were mainly divided into 4 categories in both H vs. SA (Fig. 4F) and H vs. A (Fig. 4G) groups: spliceosome, RNA transport, Lamin tail domain, and ribosome. Moreover, 69 sequence motifs were predicted from all the identified phosphorylation sites with Motif-x (Fig S1).

For the apoptosis pathway, 14 related proteins, including caspase3, were higher in GCs of SA than H. The protein levels of laminA/C and lamin B were higher in GCs of H than that of SA, however, phosphorylation levels of lamin A/C (S22 and S636) and lamin B1 (S308 and S534) were higher in GCs from SA than from H. lamins, major structural proteins of the cell, are targeted for destruction early in apoptosis (39). Lamin A/C is cleaved to a small fragment (28 kDa), specifically by caspase6, but not by other caspases, during the induction of apoptosis (39, 40). The protein-protein interaction networks also showed direct interactions between caspase6 and lamin A/C (Fig. 4 F, G). Proteomic analysis showed no difference in the caspase 6 protein levels in GCs from H, SA, and A. The phosphorylation level of caspase6 (S76) was higher in GCs from H than that from SA. Phosphorylation occurs in all structural domains of caspases typically resulting in caspase inactivation by preventing formation of the active, cleaved caspase (41). Western blot results confirmed the protein levels of caspase6 and laminA/C, as well as phosphorylation level of laminA/C at S22 in GC during follicular atresia, and showed that both cleaved caspase6 and cleaved laminA/C were higher in GC from A and SA than that from H (Fig. 5A). The expression pattern of laminA/C and caspase6 were also detected by immunofluorescence. Both laminA/C and caspase6 were expressed in the cytoplasm of GCs in H follicles. However, as the follicles underwent atresia, lamin A/C and caspase 6 entered the nucleus, diffusely expressing in the apoptotic GCs, and their staining signals are also weaker (Fig. 5B). A previous study showed that phosphomimetic substitution of S22 in lamin A/C resulted in an increase in laminA/C in the nucleoplasm (42). However, a mutation of S22 to Ala on laminA significantly inhibits laminA disassembly in mitotic cells (43). The levels of phosphorylated S636 in laminA have been shown to increase 2-fold in mitotic HeLa S3 cells (44). Activation of caspase3 can result in activation of pro-caspase6, and activation of pro-caspase6 can also result in activation of caspase3, resulting in a protease amplification cycle (45, 46). Collectively, we deduce that phosphorylation at S76 may block the activity of caspase6, therefore preventing lamin A/C from cleavage. Phosphorylation of laminA/C at S22 and S636 may contribute to laminA/C disassembly or cleavage, thus promoting apoptosis of GCs during follicular atresia (Fig. 5C).

**Fig 5.**
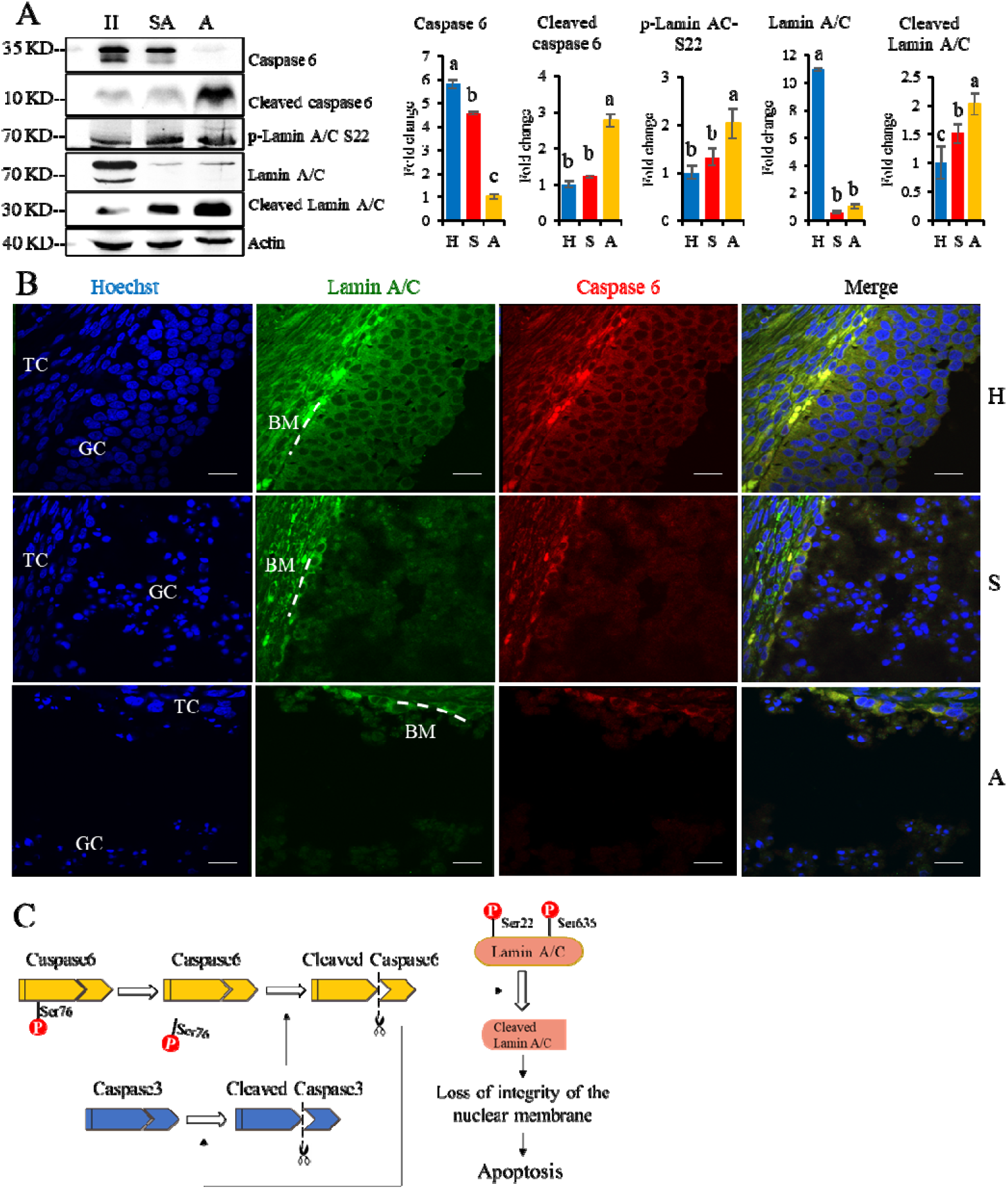
The expressions of caspase6, cleaved caspase6, p-laminA/C-S22, laminA/C, and cleaved laminA/C in GCs from healthy (H), slightly atretic (SA), and atretic (A) follicles, as well as possible regulatory pathways of caspase6, caspase3, and laminA/C. A. Protein levels of caspase6, cleaved caspase6, p-laminA/C-S22, laminA/C, and cleaved laminA/C in GCs from H, SA, and A follicles were detected by Western blotting. Quantitative analysis of these proteins is shown in histograms. The bars are labeled with completely different letters (a, b, c) indicating significant difference, *P* < 0.05. B. The expression of laminA/C and caspase6 in GCs of follicles in different health states was detected by immunofluorescence. Nuclei were stained with Hoechst. Scale bar=100 μm. TC=theca cell; GC=granulosa cell; BM=basement membrane. C. Hypothetical model of regulatory pathways of caspase6, caspase3, and laminA/C in GCs during follicular atresia. The data are representative of at least three independent experiments.

### 3.4. Transcription Factor dynamics in GCs during follicular atresia

To investigate dynamics of transcription factors along the steps of porcine follicular atresia, all 1,490 known porcine transcription factors (TFs) from AnimalTFDB3 were analyzed (24). A total of 477 TFs were identified and 359 TFs were quantified in the proteome. Cluster 1 comprised 127 TFs, which were upregulated in H compared with SA and A. Among these, MAX, YBX1, zf-C2H2, HMGB2, NR5A2, TOX2, HMG and ZMAT1 had a FC ≥ 3, indicating that these TFs may play a critical role in follicular development. Cluster 3 consisted of 102 TFs, which were upregulated in SA and A compared with H. Among these, STAT5A, MITF, ZNF260, SAND, LBX2 Homeobox, STAT5B, CBFB, XPA, and STAT1 had a FC ≥ 2, suggesting that they are important in the initiation of atresia. Cluster 6 included 46 TFs, which were upregulated in A compared with SA and H. Among these, E2F, CARHSP1, MYT1L, HMGA, and ZNF516 had a FC ≥ 2, indicating that they are vital in the late stage of atresia (Fig. 6A).

**Fig 6.**
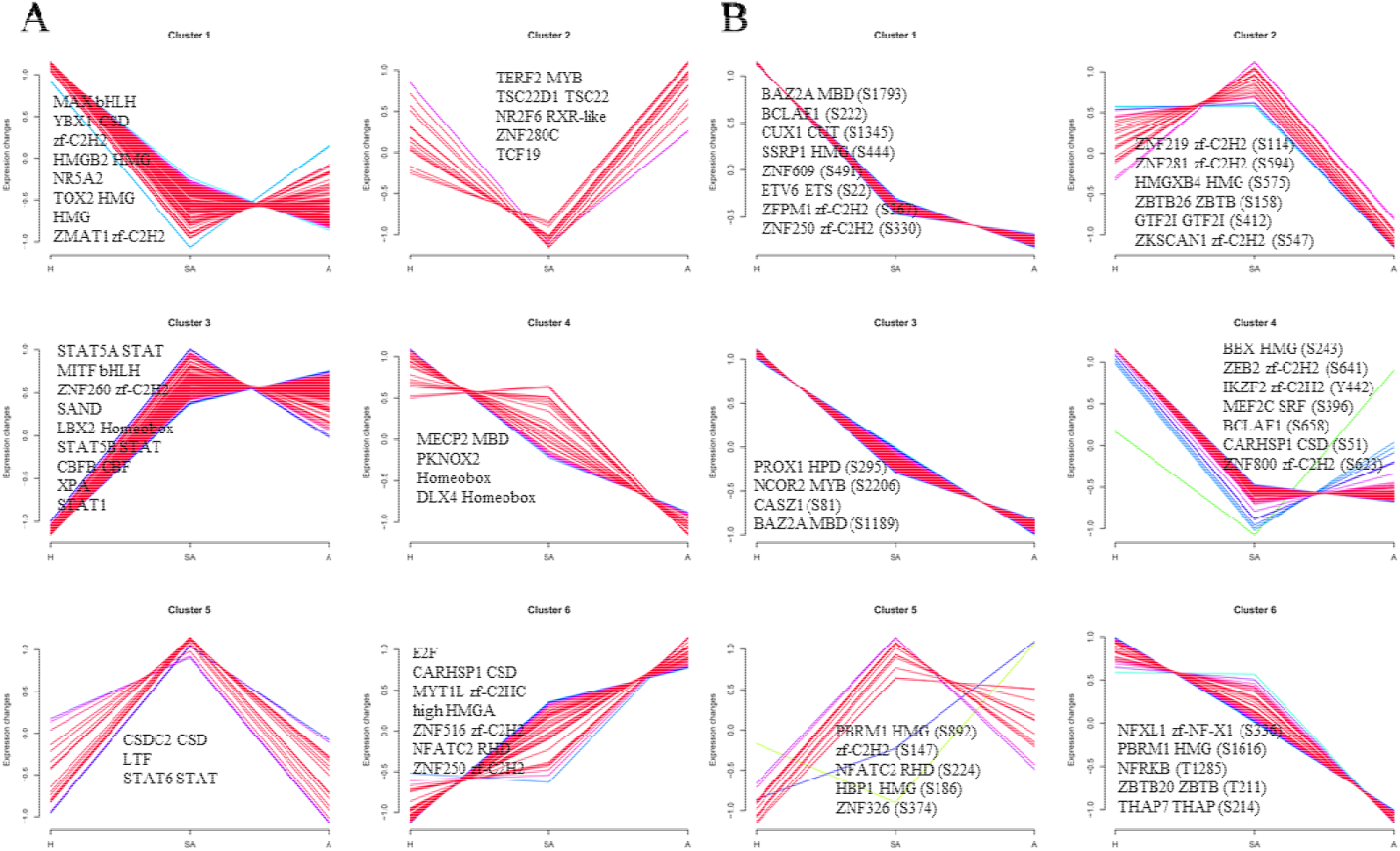
Clustering analysis of TF expression patterns and changes of phosphorylated TFs in GCs from three different follicular statuses. A. Clustering analysis of TF expression patterns in proteome. B. Clustering analysis of phosphorylated TF Phosphoproteome. Each line indicates the relative abundance of each protein. Selective proteins or phosphosites in each cluster are shown.

In total, 173 of 307 identified TFs were quantified, and 572 of 850 identified sites were quantified in the phosphoproteome. Cluster 1, cluster 3, cluster 4, and cluster 6 consisted of 419, 142, 139 and 37 phosphorylation sites, respectively. All were upregulated in H compared with SA and A. Cluster 2 included 23 phosphorylation sites, which were highest in SA, second highest in H, and lowest in A. Cluster 5 consisted 14 phosphorylation sites, which were highest in SA, second highest in A, and lowest in H (Fig. 6B).

### 3.5. Kinase signaling dynamics in GCs during follicular atresia

A total of 2,762 regulatory relationships were predicted between 238 protein kinases and 1,140 phosphorylated sites on 606 proteins identified by phosphoproteomics (S4 File). The results of predictive analysis of kinase activity showed that 27 kinases in H were strongly activated, including several cell cycle kinases, including CK2 (CSNK2A1/2), CDC7, CDK5, CDK7, CDK13, and MAPKs (MAPK1, MAPK8, MAPK9, MAPK14). However, NEK2 was blocked in H. In addition, 9 kinases in A were predicted to be inactive, which also included several cell cycle kinases, CK2, CDK2, CDK3, CLK2, and CDK11B (Fig. 7A). Kinase activity inferred from phosphoproteomic data indicated that PBK, CDK5, and CDK12 were upregulated in H/SA. In addition, AKT3, PIK3C3, DYRK1A, CDC7, and ATR were upregulated in H/A group (Fig. 7B). These upregulated kinases are mainly related to cell cycle, mitosis, and transcription. Kinase regulatory networks of PBK, CDK5, and CDK12 are shown in Fig. 7C, and networks of AKT3, PIK3C3, DYRK1A are shown in Fig. 7D. Of note, CDK12 was predicted to phosphorylate Setd2 at S493, S594, S862, and S2039. Setd2 is specific methyltransferase for lysine-36 of histone H3, and methylation of this residue is associated with active chromatin (47). PIK3C3 was predicted to phosphorylate SQSTM1 at S272 and AMBRA1 at S52. Both of these proteins are autophagy regulators (48); and phosphorylation at these two sites inhibits autophagy (49, 50). Transcription factor FOXO1 (S305 and S336) was phosphorylated by DYRK1A kinase and is involved in many biological processes including apoptosis, cell cycle arrest, stress resistance, glucose metabolism, cellular differentiation and development, and tumor suppression (51). Phosphorylation of FOXO proteins by DYRK1A has been reported to promote export of these proteins to the cytoplasm (inactive), while phosphorylation by JNK can trigger the localization of FOXO proteins from the cytoplasm to the nucleus (51). The two phosphosites of FOXO1 found in this study is important for follicular development and prevent GCs from apoptosis; and may be valuable for developing compounds that disrupt the phosphorylation of FOXO1 by DYRK1A, which could serve as potential choice for cancer-related drug design. DYRK1A was also predicted to phosphorylate Sirtuin 1, a histone deacetylase, at S26, S47, and S592. AKT3 was predicted to phosphorylate PDCD4 at S458, which may lead ubiquitination and degradation of PDCD4, therefore inhibiting GC apoptosis and promoting translation (52). The PDCD4 protein may be phosphorylated through the EGF-activated PI3K-AKT-mTOR-S6k1 signaling pathway and degraded in the proteasome system (53). AKT3 was also predicted to phosphorylate several HSP proteins, which play roles in stress resistance and actin organization.

**Fig 7.**
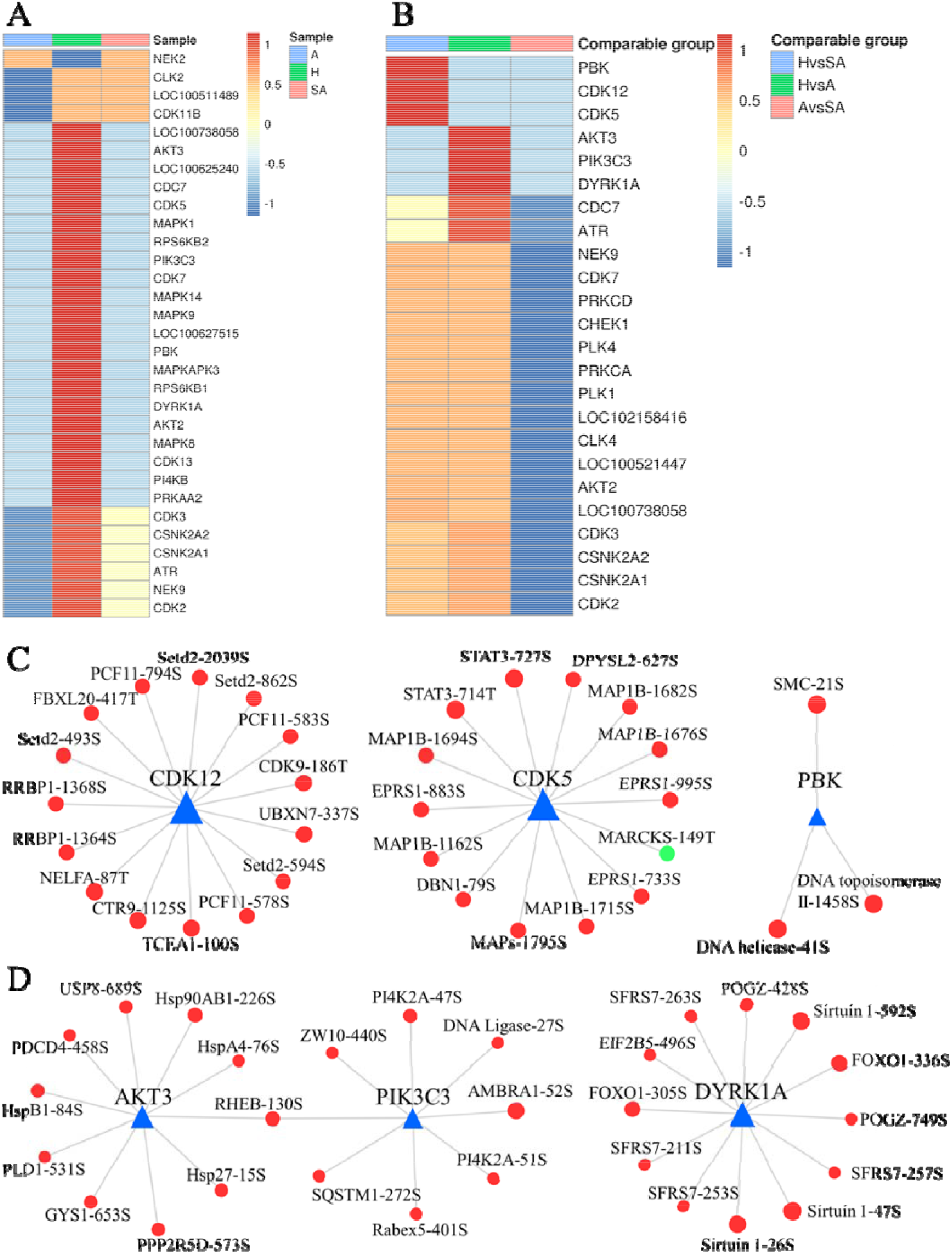
Dynamic changes of kinase activity and the key kinase-substrate regulatory network in GCs during follicular atresia. A. Heat map of kinase activity scores in H, SA, and A. Columns are sample names and rows are phosphokinases (rows are clustered and Z-score normalized). B. Heat map of kinase activity matrix in comparable groups. The columns are the comparison group names and the rows are phosphokinases (rows are clustered and Z-score normalized). C. Map of key kinase-substrate regulatory networks in H/SA. D. Map of key kinase-substrate regulatory networks in H/A.

## Discussion

Follicular atresia is a complicated process that limits the potential fertility of domestic animals. In this study, comprehensive proteomic and phosphoproteomic analyses were applied in GCs from different quality categories of follicles. Transcriptome analyses in GCs during follicular development and atresia have been performed in porcine (18, 19, 54), sheep (55), bovine (56), and humans (57). Some key genes and pathways were found to be related to these processes. Proteomic analysis had also conducted in porcine follicular fluid during follicle development. The patterns of protein-to-protein interactions throughout follicle development, stage-specific proteins, and major pathways have been uncovered (58). However, the present study is the first proteomic and phosphoproteomic analysis of the process of mammalian follicular atresia. The large number of DEPs and DEPPs may provide valuable clues for future functional studies. The changes of TFs and kinases in GCs during atresia were also unveiled. An in-depth understanding of the mechanism of follicular atresia is helpful for the advancement of reproductive technologies, such as three-dimensional follicle culture, and developing artificial ovaries, and may also provide some new ideas for cancer treatment.

In human assisted reproduction and livestock production, it is critical to distinguish between healthy and atretic follicles and to evaluate oocyte quality. Morphological changes, apoptosis of GCs, detachment of the GC layer from the follicular BM, and an increased ratio of progesterone to estrogen (E_2_) can serve as powerful evidence of follicular atresia (16, 20, 21). Evaluation of oocyte quality mainly depends on morphologic characteristics (59). In this study, some DEPs (MIF, beta catenin) and DEPPs (S22 of laminA/C, S76 of caspase6) could serve as potential novel biomarkers to distinguish follicles of different health status, and to evaluate oocyte quality.

Some novel phosphorylation sites of key proteins were found in the phosphoproteomic analysis, which may have important significance in the functional study of corresponding proteins and in clinical applications. For example, regulation of caspase6 activation is not clear. In this study, the phosphorylation level of S76 of caspase6 (same as S79 of caspase6 in human) was found to be higher in GCs from H compared with SA or A, which may be a key phosphosite to suppress caspase6 activation. Increasing evidence has shown that caspase6 is highly involved in axon degeneration and neurodegenerative diseases, such as Huntington’s disease and Alzheimer’s disease (60). Inhibiting the function of caspase6 may have therapeutic potential for various neurodegenerative disorders (60, 61). In addition, five phosphorylation sites (S30, S269, S274, S275, and T319) of HSD17B1 had higher levels of phosphorylation in GCs from H than from SA and A, which may contribute to the catalytic activity of HSD71B1 and promote E_2_ synthesis. This may explain the change in the ratio of progesterone to E_2_ during atresia. HSD17B1 is overexpressed in E_2_-dependent cancers, such as breast, endometrial, and ovarian cancer (62, 63). Therefore, these five phosphosites may provide new insights into the treatment of these types of cancers. Another example is the change of phosphorylation level of the gap junction alpha-1 protein (Connexin43) at S328 and S262, which may affect gap junction formation and permeability during mitosis, thus affecting the transfer of materials between GCs (64, 65).

DEPs and DEPPs in biological process were mainly concentrated in cellular process, metabolic process, single-organism process, biological regulation, response to stimulus, localization, and signaling. Results of the current study revealed that blood coagulation, proteolysis, and oxoacid metabolic process were elevated in SA, while peptide metabolic process, amide biosynthetic process, and chromatin organization process were damaged in SA. Nucleobase-containing small molecule metabolic process and actin cytoskeleton organization were mapped to be higher in A; however, cell redox homeostasis was destroyed in A. These most significantly affected biological processes may better explain the mechanisms of atresia.

Apoptosis, proteasome, and complement and coagulation cascades pathways were activated in SA, which may play critical roles in the initiation of atresia. Regulation of the actin cytoskeleton, NF-kappa B signaling pathway, and aminoacyl-tRNA biosynthesis pathway were increased in A, indicating that these pathways may contribute to the final degeneration of follicles. However, peroxisome, glycolysis/gluconeogenesis, vasopressin-regulated water reabsorption, mismatch repair, galactose metabolism, gap junction pathway, and steroid hormone biosynthesis pathways were upregulated in H, which may contribute to follicular development, while their downregulation maybe the main reason of follicular atresia. Predicted pathways and the corresponding DEPs and DEPPs will be valuable for conducting further investigations and verification analysis of porcine follicular atresia. The proteome and phosphoproteimic analysis in the current study found some of the same KEGG pathways as those found in the previous transcriptome: apoptosis, AGE-RAGE signaling pathway in diabetic complications, fluid shear stress and atherosclerosis, ECM–receptor interaction, and Ribosome (19). Our analysis also revealed some new KEGG pathways that are involved in atresia, including peroxisome, cysteine and methionine metabolism, glycolysis/gluconeogenesis, ubiquitin mediated proteolysis, spliceosome, mismatch repair, galactose metabolism, regulation of actin cytoskeleton, NF-κB signaling pathway, proteasome, and aminoacyl-tRNA biosynthesis.

In the regulation of actin cytoskeleton pathway, the levels of 23 proteins were elevated in A compared to H. Among these, cofilin-1 and integrin β2 were the most variable proteins. The actin cytoskeleton acquires a new conformation; a sphere-like structure which separates the apoptotic blebs from the main bulk of the cell, and plays an important role in transmitting survival signals exerted by the BM, laminin and growth factors which activate tyrosine kinase receptors (15). Active cofilin can regulate the rearrangement of the actin cytoskeleton and required to initiate progesterone secretion by preovulatory GCs via the LHR-PKA signaling pathway (66). Integrins are the major cellular receptors mediating adhesion to the ECM, and integrin signaling regulates actin reorganization, cell proliferation, apoptosis, gene expression, differentiation, and cell migration (67). During folliculogenesis, the expression of integrin subunits in GCs varies significantly between animal species (67). It has been reported in sheep that α6 associates with the β1 integrin subunit to form the α6β1 laminin receptor in GCs of healthy follicles, and the expressions of both subunits decrease with atresia (68). Our data showed that integrin β2 in porcine GCs was about 4 times higher in A than H, which may be important for GC apoptosis and follicular atresia.

In the NF-kappa B signaling pathway, the levels of 9 proteins were increased in A compared with H. Among these, Bcl10 was the most variable protein, which contains a caspase recruitment domain (CARD), and has been shown to induce apoptosis and to activate NF-κB (69). TNF receptor-associated factor 6, which functions as a signal transducer in the NF-kappa B pathway that activates I kappa B kinase (IKK) in response to proinflammatory cytokines (70, 71), and was also elevated in GCs from A follicles. Our earlier study showed that inhibition of NF-κB increased autophagy via JNK signaling, and promoted progesterone secretion in porcine GCs (72).

In the aminoacyl-tRNA biosynthesis pathway, the levels of 13 proteins, including seryl-tRNA synthetase, leucyl-tRNA synthetase, tyrosyl-tRNA synthetase, threonyl-tRNA synthetase, prolyl-tRNA synthetase, and methionyl-tRNA synthetase were higher in A than in H, indicating that the aminoacyl-tRNA biosynthesis or metabolism was increased in GCs from A. ARSs play a vital role in protein synthesis by linking amino acids to their cognate transfer RNAs (tRNAs). Aminoacyl-tRNA has also been found to have functions in several other biosynthetic pathways, such as lipid, and protein degradation (73). The role of the aminoacyl-tRNA biosynthesis pathway in atretic follicles needs to be further studied.

In the ECM-receptor interaction, 8 proteins mapped to be higher in GCs from H than from SA. Among these, dystroglycan had the largest FC, and was 4.27 time higher in GCs from H than that from A. ECM plays an active and complex role in regulating the morphogenesis of cells that contact it, influencing their survival, migration, proliferation, and metabolic functions (74). In the mammary epithelium, loss of cell adhesion to the BM due to ECM proteolysis is believed to be sufficient to initiate apoptosis and involution of the gland (75). Growth factors (such as the basic fibroblast growth factor) and/or adhesion molecules within the ECM are also related to cell survival, as well as proliferation (76, 77). Increasing evidence suggests that various ovarian ECM components present in the follicular BM, around GCs, and in the follicular fluid are important regulators of GC activity (67). ECM can regulate the expression of StAR protein and P450scc in rat GCs, which stimulates progesterone production. Rat GCs cultured in ECM are protected from apoptosis and have a well-developed actin cytoskeleton that is extensively spread out (74). An earlier study suggested that the inner layers of GCs, distal to the BM, are more sensitive to apoptotic signals than cells in close contact with the BM where the antiapoptotic signal of the BM, and in particular of laminin, has a direct effect (74). In this study, laminin, collagen, integrin α6 and proteoglycan βDG were higher in GCs from H than that from SA. Change of integrin α6 in porcine GCs was consistent with that in GCs of healthy follicles in sheep (68).

In the ubiquitin mediated proteolysis, 23 phosphosites of 13 proteins were higher in GCs from H than SA. Phosphosites of S100, S313, Y1052, and S1053 in E3 ubiquitin-protein ligase TRIP12, as well as S841 in E2 ubiquitin-conjugating enzyme UBE2O had a FC ≥ 3. Numerous cellular processes regulated by ubiquitin-mediated proteolysis include the cell cycle, DNA repair and transcription, protein quality control and the immune response. Our recent study showed that melatonin may maintain follicular health by inducing BimEL ubiquitination to inhibit the apoptosis of GCs (78).

Using the TF database, we found some key TFs and phosphorylated TFs were found, which are involved in normal development of follicles, initiation of follicle atresia, and eventually degeneration of follicles. For example, BCLAF1 is a transcriptional repressor that interacts with several members of the BCL-2 family of proteins and overexpression of this protein induces apoptosis (79). BCLAF1 is an important NF-κB signaling transducer and C/EBPβ regulator in DNA damage-induced senescence (80). Our data indicate that the phosphorylation levels of BCLAF1 at S222 and S658 were higher in SA and A compared with H, which may contribute to its ability to induce apoptosis.

Kinase analysis revealed that there are more active kinases in GCs from H than from SA and A. These predicted differential kinases are mainly involved in cell cycle, mitosis, and transcription. The regulation networks of these key kinases were established. The inactivated kinases in GCs from A may contribute to the atresia process. TFs such as STAT3 (T714 and S727), FOXO1 (S305 and S336), RUNX1 (T22 and S220), SP1 (S59), TCEA1 (S100), and UBTF (S389) were predicted to be regulated by these kinases, which may better explain the mechanism of follicular atresia.

## Conclusion

In summary, this study provides a comprehensive profile of DEPs and DEPPs in healthy, slightly atretic, and atretic antral follicles in the porcine. A large number of proteins and phosphosites were found to be different in the GCs from three different quality categories of follicles. Some potential novel biomarkers for follicular atresia, such as MIF, were found. Some novel phosphosites of key proteins were found in the phosphoproteomic analysis, which may have significance in functional studies of corresponding proteins and in clinical applications. Further analysis of the DEPs and DEPPs revealed several core proteins, key phosphosites, biological processes, KEGG pathways, TFs, and kinases that are involved in atresia. The results in the current study could lead to a better understanding of the molecular regulation of ovarian follicular atresia.

This article contains supplemental data.

## Abbreviations

GC: granulosa cell
H: healthy follicles
SA: slightly atretic follicles
A: atretic follicles
FC: fold change
DEPs: differential expressed proteins
DEPPs: differential expressed phosphorylated proteins
TF: transcription factor
PTM: posttranslational modification
BM: basement membrane
GO: Gene Ontology
KEGG: Kyoto Encyclopedia of Genes and Genomes
TC: theca cell
ECM: extracellular matrix

## Data Availability

Data are available via ProteomeXchange with identifier PXD020899.

## Acknowledgement

This work was supported by grants from the National Key R&D Program of China (2017YFC1002003) and National Natural Science Foundation of China (31672419, 31470077). We thank Jingjie PTM Biolab for the excellent technical assistance and mass spectrometric analysis. We thank LetPub (www.letpub.com) for its linguistic assistance during the preparation of this manuscript.

## Author contributions

F.Y., Q.L., Y.C., H.Y, and H.W. performed research; F.Y. and Q.L. analyzed data; F.Y. and Q.L. wrote the paper; F.Y., Q.L. and S.Z. designed research.

