## Supplementary material for "Integrative proteomic and phosphoproteomic analysis of granulosa cells during follicular atresia in porcine": Fig S1--motif-x--Phosphoproteomic.pdf

| Motif Logo | Motif | Motif Score | Foreground Matches | Foreground Size | Background Matches | Background Size | Fold Increase |
|---|---|---|---|---|---|---|---|
| 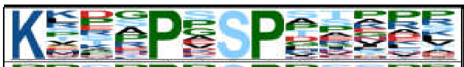 | K...P.SP..... | 38.15 | 42 | 5709 | 218 | 733195 | 24.74 |
| 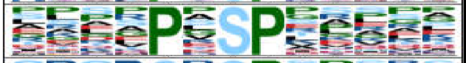 | .....P.SP..... | 32 | 483 | 5667 | 6078 | 732977 | 10.28 |
| 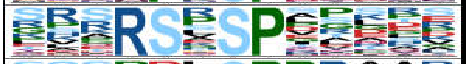 | ...RS.SP..... | 39.51 | 109 | 5184 | 551 | 726899 | 27.74 |
| 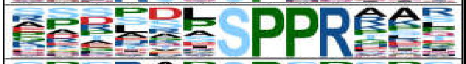 | ......SPPR... | 41.54 | 55 | 5075 | 280 | 726348 | 28.11 |
| 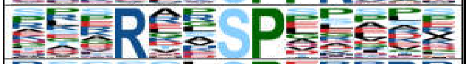 | ...R..SP..... | 32 | 252 | 5020 | 2472 | 726068 | 14.74 |
| 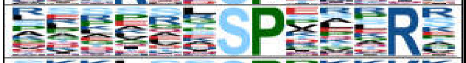 | ......SP...R. | 32 | 215 | 4768 | 2662 | 723596 | 12.26 |
| 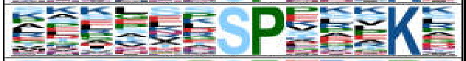 | ......SP...K. | 32 | 186 | 4553 | 2147 | 720934 | 13.72 |
| 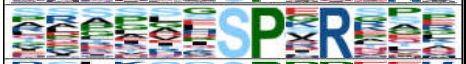 | ......SP.R... | 32 | 158 | 4367 | 1879 | 718787 | 13.84 |
| 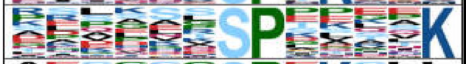 | ......SP....K | 31.18 | 133 | 4209 | 1681 | 716908 | 13.48 |
| 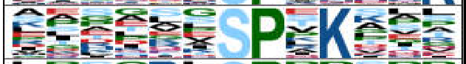 | ......SP.K... | 29.25 | 118 | 4076 | 1591 | 715227 | 13.01 |
| 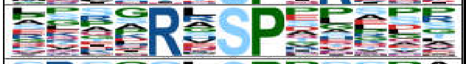 | ....R.SP..... | 27.82 | 135 | 3958 | 2138 | 713636 | 11.38 |
| 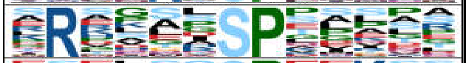 | .R....SP..... | 27.91 | 112 | 3823 | 1745 | 711498 | 11.95 |
| 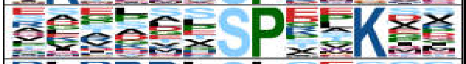 | ......SP..K.. | 25.94 | 82 | 3711 | 1287 | 709753 | 12.19 |
| 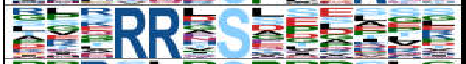 | ...RR.S...... | 32 | 196 | 3629 | 3194 | 708466 | 11.98 |
| 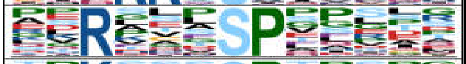 | ..R...SP..... | 23.64 | 80 | 3433 | 1478 | 705272 | 11.12 |
| 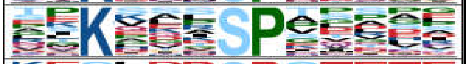 | ..K...SP..... | 23.86 | 68 | 3353 | 1221 | 703794 | 11.69 |
| 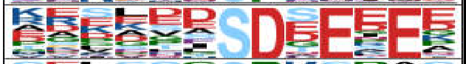 | ......SD.E.E. | 40.06 | 55 | 3285 | 436 | 702573 | 26.98 |
| 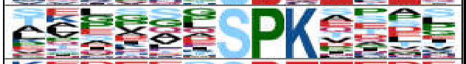 | ......SPK.... | 24.04 | 57 | 3230 | 981 | 702137 | 12.63 |
| 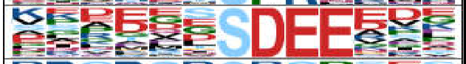 | ......SDEE... | 43.3 | 39 | 3173 | 365 | 701156 | 23.61 |
| 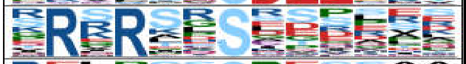 | .R.R..S...... | 32 | 139 | 3134 | 3269 | 700791 | 9.51 |
| 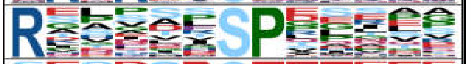 | R.....SP..... | 24 | 66 | 2995 | 1291 | 697522 | 11.91 |
| 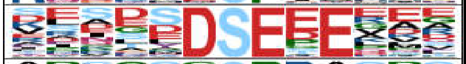 | .....DSE.E... | 42.23 | 40 | 2929 | 445 | 696231 | 21.37 |
| 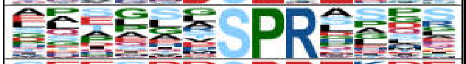 | ......SPR.... | 22.7 | 59 | 2889 | 1274 | 695786 | 11.15 |
| 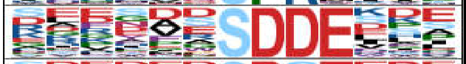 | ......SDDE... | 38.36 | 27 | 2830 | 244 | 694512 | 27.16 |
| 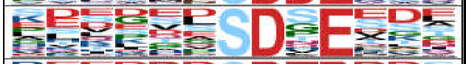 | ......SD.E... | 32 | 76 | 2803 | 2412 | 694268 | 7.8 |
| 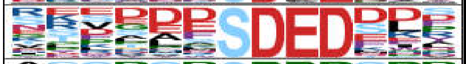 | ......SDED... | 40.86 | 30 | 2727 | 285 | 691856 | 26.71 |
| 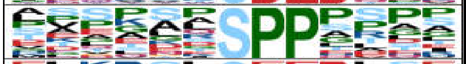 | ......SPP.... | 23.18 | 87 | 2697 | 2279 | 691571 | 9.79 |
| 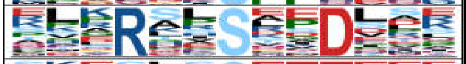 | ...R..S..D... | 30.88 | 69 | 2610 | 1599 | 689292 | 11.4 |
| 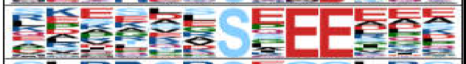 | ......S.EE... | 28.81 | 110 | 2541 | 5115 | 687693 | 5.82 |
| 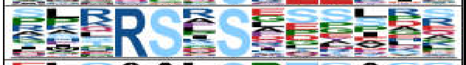 | ...RS.S...... | 32 | 80 | 2431 | 3638 | 682578 | 6.17 |
| 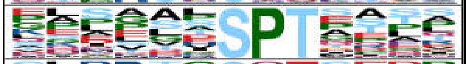 | ......SPT.... | 23.9 | 59 | 2351 | 1401 | 678940 | 12.16 |
| 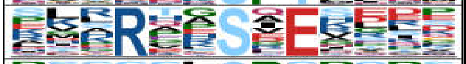 | ...R..S.E.... | 25.15 | 45 | 2292 | 1830 | 677539 | 7.27 |
| 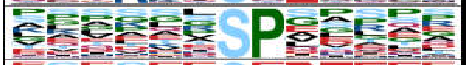 | ......SP..... | 16 | 337 | 2247 | 19312 | 675709 | 5.25 |
| 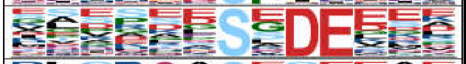 | ......S.DE... | 32 | 69 | 1910 | 2280 | 656397 | 10.4 |
| 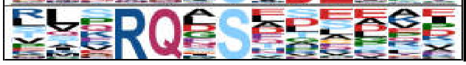 | ...RQ.S...... | 24.2 | 39 | 1841 | 1305 | 654117 | 10.62 |

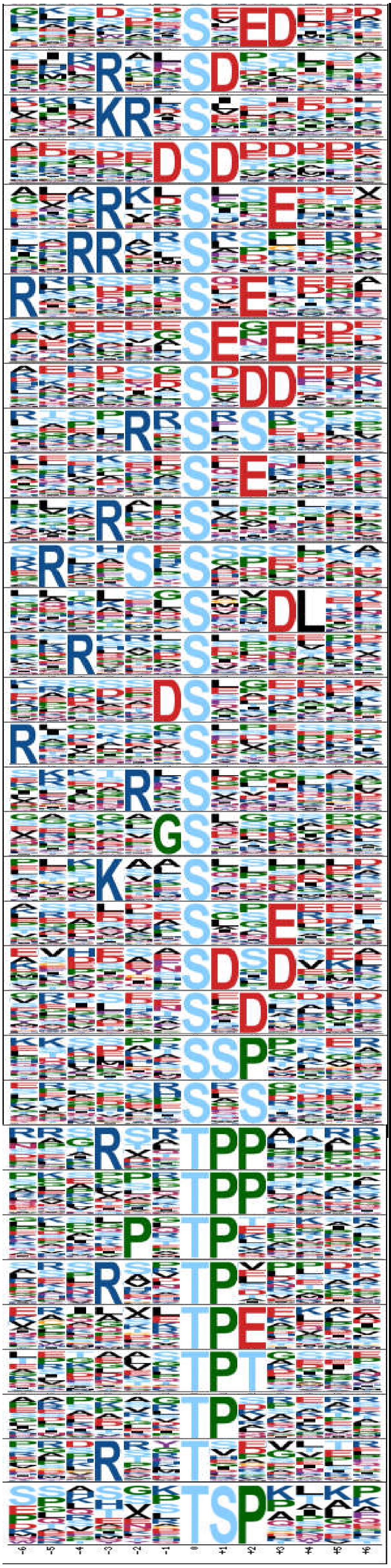

| | | | | | | |
|---|---|---|---|---|---|---|
| ......S.ED... | 27.06 | 55 | 1802 | 2794 | 652812 | 7.13 |
| ...R..SD..... | 22.5 | 29 | 1747 | 1043 | 650018 | 10.35 |
| ...KR.S...... | 29.49 | 49 | 1718 | 2131 | 648975 | 8.69 |
| .....DSD..... | 32 | 50 | 1669 | 1857 | 646844 | 10.44 |
| ...R..S..E.... | 23.57 | 29 | 1619 | 1595 | 644987 | 7.24 |
| ..RR..S...... | 22.96 | 28 | 1590 | 1620 | 643392 | 6.99 |
| R......S.E.... | 22.69 | 29 | 1562 | 1826 | 641772 | 6.53 |
| ......SE.E... | 22.82 | 37 | 1533 | 2929 | 639946 | 5.27 |
| ......S.DD... | 29.26 | 42 | 1496 | 1792 | 637017 | 9.98 |
| ....R.S.S.... | 23.05 | 42 | 1454 | 3258 | 635225 | 5.63 |
| ......S.E.... | 16 | 159 | 1412 | 30848 | 631967 | 2.31 |
| ...R..S...... | 16 | 149 | 1253 | 20340 | 601119 | 3.51 |
| .R..S.S...... | 22.51 | 38 | 1104 | 3713 | 580779 | 5.38 |
| ......S..DL.. | 22.7 | 36 | 1066 | 2942 | 577066 | 6.62 |
| ..R...S...... | 15.95 | 123 | 1030 | 30479 | 574124 | 2.25 |
| .....DS...... | 13.08 | 97 | 907 | 25397 | 543645 | 2.29 |
| R.....S...... | 12.85 | 96 | 810 | 27029 | 518248 | 2.27 |
| ....R.S...... | 9.92 | 69 | 714 | 20268 | 491219 | 2.34 |
| .....GS...... | 10.27 | 97 | 645 | 35179 | 470951 | 2.01 |
| ...K..S...... | 8.63 | 69 | 548 | 25407 | 435772 | 2.16 |
| ......S..E... | 7.23 | 57 | 479 | 22623 | 410365 | 2.16 |
| ......SD.D... | 14.06 | 14 | 422 | 1064 | 387742 | 12.09 |
| ......S.D.... | 7.52 | 47 | 408 | 18387 | 386678 | 2.42 |
| ......SSP.... | 12.6 | 23 | 361 | 3961 | 368291 | 5.92 |
| ......S.S.... | 6.11 | 72 | 338 | 43491 | 364330 | 1.78 |
| ...R..TPP.... | 38.33 | 26 | 651 | 167 | 459774 | 109.96 |
| ......TPP.... | 32 | 113 | 625 | 2512 | 459607 | 33.08 |
| ....P.TP..... | 28.86 | 73 | 512 | 3179 | 457095 | 20.5 |
| ...R..TP..... | 27.53 | 39 | 439 | 1432 | 453916 | 28.16 |
| ......TPE.... | 22.02 | 33 | 400 | 2088 | 452484 | 17.88 |
| ......TPT.... | 22.69 | 26 | 367 | 1528 | 450396 | 20.88 |
| ......TP..... | 16 | 95 | 341 | 20538 | 448868 | 6.09 |
| ...R..T...... | 10.61 | 42 | 246 | 22708 | 428330 | 3.22 |
| ......TSP.... | 19.01 | 21 | 204 | 2883 | 405622 | 14.48 |
